# Stabilising effect of modularity in antagonistic networks depends on intraguild interactions

**DOI:** 10.64898/2026.07.17.739155

**Authors:** Rémi Legrand Duchesne, Franziska Koch, Bismark Ofosu-Bamfo, Korinna T. Allhoff

## Abstract

Existing literature on ecological networks provides valuable insights into the structure-stability relation of antagonistic, mutualistic or competitive systems, but it remains unclear whether these insights also apply to networks that contain a mix of different interaction types. Here, we study the effect of modularity on stability in systems that contain not only antagonistic interactions between two guilds, but also competition, facilitation or even antagonism within each guild, inspired by Ghanaian tree-liana interaction networks. We represent these systems as structured community matrices with random interaction strengths, in which we vary both the modularity within the antagonistic subnetwork and the type of intraguild interactions. Using the eigenvalues of the community matrix to assess stability, we find that modularity in antagonistic interactions generally has a stabilising effect, in line with results on single-interaction type networks. We furthermore find that the magnitude of this effect is modulated by the type of intra-guild interaction under consideration and is largest when these interactions describe facilitation. We explain these findings via a shift in the balance between self-reinforcing and self-damping feedback loops. Our results highlight the need to study how patterns in inter-and intraguild interactions jointly affect ecosystem stability.

## 1 Introduction

The relationship between the structure of species interaction networks and their stability has been the subject of extensive research in theoretical ecology. Classical studies based on random matrix theory demonstrate that network stability depends on species richness, mean interaction strength, connectance, as well as on the type of interaction under consideration (Allesina and Tang, 2012, 2015; May, 1972). More precisely, networks with antagonistic interactions (e.g., food webs) are generally assumed to be more stable than those with mutualistic interactions (e.g., pollination webs) (Allesina and Tang, 2012). In antagonistic networks, stability is further enhanced by weakly connected and modular structures (Stouffer and Bascompte, 2011; Thébault and Fontaine, 2010), whereas mutualistic networks are stabilised by nested and highly connected structures (Okuyama and Holland, 2008; Rohr et al., 2014; Thébault and Fontaine, 2010). Recently, structure-stability relations have also been investigated in competitive communities, pointing to hierarchy as another stabilising structural property, as the hierarchical arrangement of links within a network mitigates network instability by keeping both short and long feedback loops relatively weak (Koch et al., 2026, 2023). These theoretical results demonstrate that structure-stability relations are not universal, as the stabilising effect of a given structural pattern within a network depends heavily on the type of network under consideration.

While theoretical studies often focus on networks with only a single type of interaction, species in real systems naturally engage in multiple interaction types (Fontaine et al. (2011); Kéfi et al. (2017)). For example, plant species simultaneously interact with herbivores (antagonism) and pollinators (mutualism) (Jäger et al., 2026; Melián et al., 2009) and even when considering only plant-plant interactions, one can find both facilitation and competition, depending on plant density and/or environmental conditions (Bimler et al., 2024; Holmgren et al., 1997; Zhang and Tielbörger, 2020). Structure-stability relations in complex interaction networks with multiple interaction types have received increasing attention in the recent literature (see for example Domínguez-García and Kéfi (2024); Kéfi et al. (2016); Yan (2022)). For instance, Yan (2022) demonstrated that the effects of subnetwork nestedness on local stability and persistence of multilayer networks were highly dependent on interconnection pattern and also on subnetwork complexity. Nevertheless, the conditions under which existing theoretical insights into structure-stability relations from single-interaction type networks also hold for realistic networks that contain a mix of interaction types remains to be fully explored, and this is in particular true for the stabilising effect of modularity in predominantly antagonistic networks.

Here, we use Ghanaian tree-liana interaction networks as inspiration to investigate structure-stability relations of systems with multiple interaction types. The interaction between trees and lianas itself is considered antagonistic, given that lianas depend on tree support to have access to the light-rich canopy (Burnham, 2002) and that they have a direct negative effect on trees via reduced tree growth and fecundity, as well as increased tree mortality (Estrada-Villegas et al., 2022; Estrada-Villegas and Schnitzer, 2018; García León et al., 2018; Meunier, 2026; Pérez-Salicrup, 2001; Schnitzer and Bongers, 2002; Schnitzer et al., 2005; Stevens, 1987; Stewart and Schnitzer, 2017). At first glance, tree-liana interaction networks therefore resemble food webs or plant-herbivore networks, suggesting that tree-liana systems that are more modular and/or weakly connected might be more stable than others (Stouffer and Bascompte, 2011; Thébault and Fontaine, 2010). Interestingly, empirical data from Ghanaian tree-liana systems indeed reveal sparsely connected networks with a relatively high degree of modularity (Addo-Fordjour et al., 2021, 2023; Ofosu-Bamfo et al., 2022). However, trees also compete for several resources, such as light availability and nutrients, and the same is true for lianas (Schnitzer et al., 2005; van Breugel et al., 2012). Additionally, lianas can facilitate each other, e.g. with new lianas using already existing lianas to climb trees (Ofosu-Bamfo et al., 2019; Pérez-Salicrup et al., 2001). In tree-liana interaction networks, antagonistic interactions between trees and lianas are thus embedded in a network together with different types of intraguild interactions, as visualised in the simplified tree-liana system shown in Fig. 1. This observed mix of interaction types, together with the relatively easy acquisition of empirical networks via co-occurrence data (Ofosu-Bamfo et al., 2022), makes tree-liana networks a suitable study system for gaining a deeper understanding of structure-stability relations in networks with multiple interaction types.

**Figure 1:**
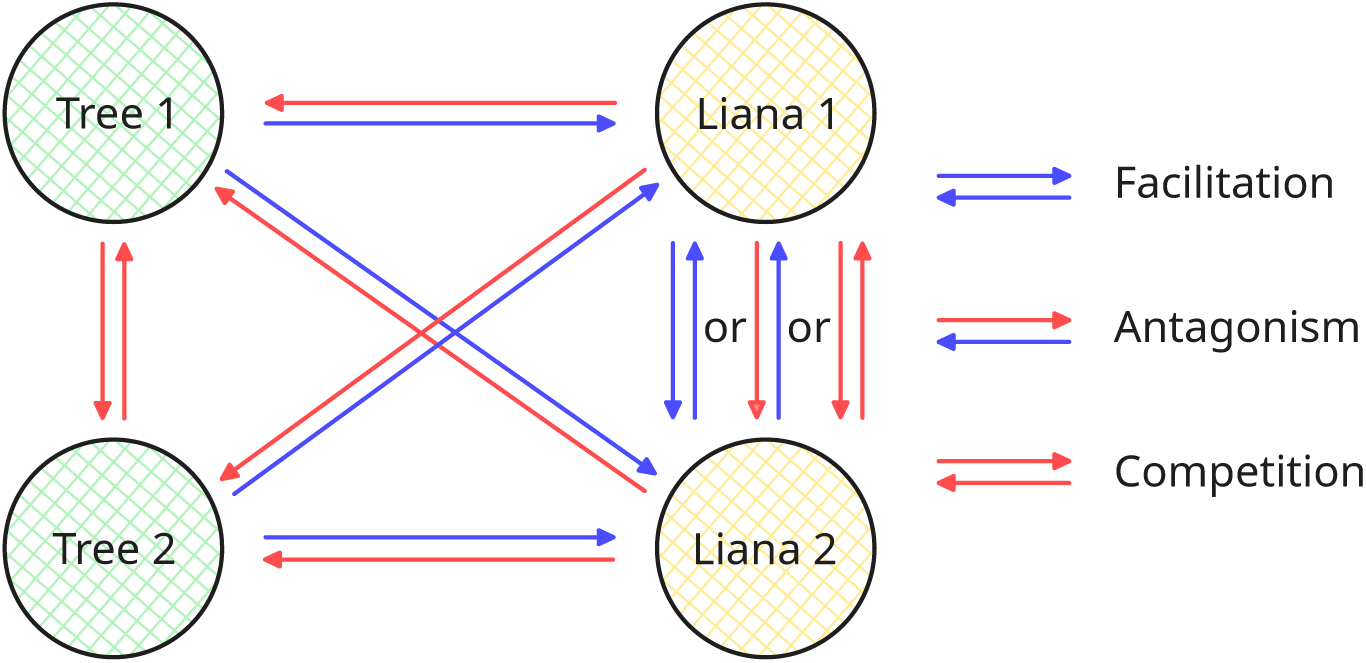
**Tree-liana systems contain various interaction types**. Interactions between trees and lianas are antagonistic: they combine a positive effect (blue arrows) of trees on lianas and a negative effect of lianas on trees (red arrows). In addition to these antagonistic interactions the network contains intraguild interactions between different tree species and between different liana species. We assume that trees always compete with each other while liana-liana interactions can be either facilitative (+/+), competitive (-/-) or antagonistic (+/-).

To study the relationship between modularity, different types of intraguild interactions and stability in tree-liana systems, we construct structured community matrices with random interaction strengths. In contrast to the matrices used in classical random matrix theory (Allesina and Tang, 2015; May, 1972), our matrices contain specific arrangements of matrix elements that are inspired by empirical observations of tree-liana systems (Addo-Fordjour et al., 2021, 2023; Ofosu-Bamfo et al., 2022). We evaluate the stability of those structured community matrices via the real part of its dominant eigenvalue and we analyse how varying the degree of modularity in the antagonistic interactions as well as the type of intraguild interactions affect positive and negative feedback loops in the systems, following previous work on trophic (Neutel et al., 2002, 2007; Neutel and Thorne, 2014) and competitive (Koch et al., 2026, 2023) networks. Feedback loops are closed chains of effects that are directly linked to system stability Levins (1974). While positive feedback loops amplify perturbations, negative feedback loops have a dampening effects. Analysing shifts in feedback loop patterns thus provides an intuitive understanding of how variation in structure leads to variation in stability. In summary, this allows us to test (1) whether modularity has a stabilising effect in systems that combine various types of biotic interactions and (2) how the type of intraguild interactions within the system modulates the relationship between modularity and stability.

## 2 Methods

We use community matrices to explore the relation between network modularity and stability in systems with different types of intraguild interactions. In line with classical approaches in theoretical ecology, we assume that our matrices represent the Jacobian matrices of an underlying (potentially non-linear) system of differential equations, evaluated at an equilibrium point (Allesina and Tang, 2012, 2015; May, 1972). Matrix elements *a_ij_* thus represent per-capita interaction strengths at this equilibrium point, which corresponds to the effect of a change in biomass of species *i* on species *j*. The sign of the matrix element corresponds to the interaction type, and its magnitude to the interaction strength. System stability is measured via the real part of the dominant eigenvalue, *Re*(*λ_d_*). A system is considered stable (meaning that it can return to its steady state after an infinitesimally small disturbance) if *Re*(*λ_d_*) *<* 0 and unstable if *Re*(*λ_d_*) *>* 0. The magnitude of *Re*(*λ_d_*) corresponds to the rate at which the system returns or moves away from steady state, so that systems with more negative *Re*(*λ_d_*) can be seen as more stable and those with more positive *Re*(*λ_d_*) can be considered more unstable.

### 2.1 Constructing structured community matrices

Our theoretical community matrices and their internal architecture are inspired by tree-liana networks (Addo-Fordjour et al., 2021, 2023; Ofosu-Bamfo et al., 2022), which is why in the following we use trees and lianas to motivate our modelling framework. Note, however, that these matrices could, in principle, represent any system with intraguild interaction in addition to antagonistic interactions between two guilds. Each community matrix is represented as a square matrix *M* of dimension (*T* + *L*) × (*T* + *L*), where *T* and *L* are the number of tree and liana species. For simplicity, we consider only systems with equal numbers of tree and liana species (*T* = *L*), which is consistent with empirical data showing that the number of tree and liana species is in the same order of magnitude (Ofosu-Bamfo et al., 2022). The matrix *M* consists of four sub-matrices, as illustrated in Fig. 2:

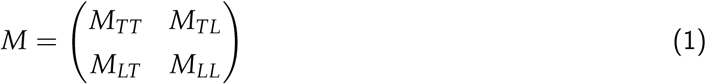

**Figure 2:**
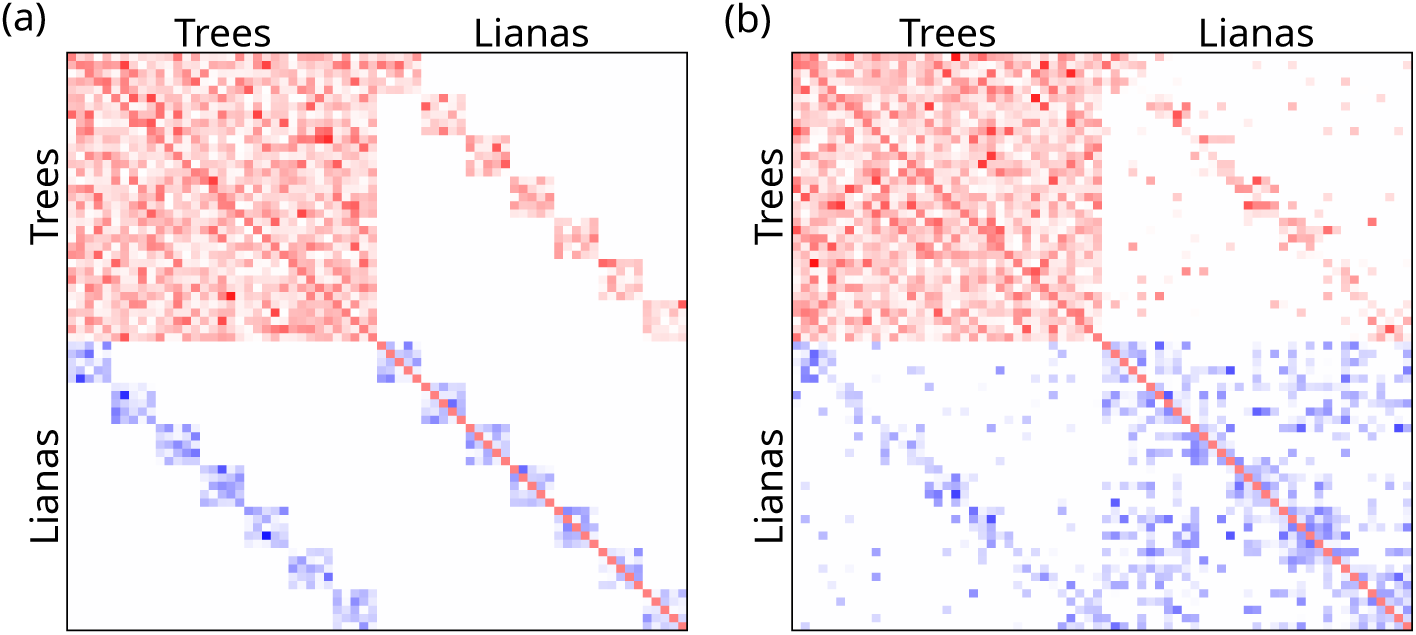
Structured community matrices with high modularity (a) and intermediate(b) modularity in the antagonistic sub-matrices. Matrix elements in the top-left represent tree-tree interactions, elements in the top-right and bottom-left represent tree-liana interactions and elements in the bottom-right represent liana-liana interactions. Red cells indicate negative interaction strengths, whereas blue cells indicate positive interaction strengths. Both matrices have been generated using the standard parameter set displayed in Table 1 but differ in their degree of modularity. a: *m* = 1, b: *m* = 0.6.

**Table 1:** Summary of model parameters. . Standard values for *C*, *n_m_*and *m* are based on the metrics from Ofosu-Bamfo et al. (2019).

| Parameter | Comments | Standard value |
| --- | --- | --- |
| Number of (tree or liana) species | We choose the same number of tree and liana species. | $n_{species} = 35$ |
| Connectance | Defined as the number of links in the antagonistic sub-matrices divided by the number of possible links in the antagonistic sub-matrices. | $C = 0.14$ |
| Number of modules | We keep the number of species fixed and assume that modules within a network have equal size. The number of modules thus also defines the size of the modules. | $n_m = 7$ |
| Input modularity | Controls the level of modularity in our community matrices. Defined as a scaling factor that describes how many links are redistributed from outside predefined modules to within. | $m \in [0.1; 1]$ |
| Scale of interaction strength distribution | For all sub-matrices, we choose normalised interspecific interaction strengths from a half-normal distribution. Larger values of the scale parameter translate into broader distributions, that is higher average interaction strengths and also higher variation in interaction strengths. | $\sigma = 0.5$ |

The first sub-matrix, *M_TT_*, captures competitive interactions between tree species for above-ground and below-ground resources (van Breugel et al., 2012) and thus contains only negative interaction strengths. We assume no spatial structure, and that tree species are well mixed in space; thus, all tree species compete with each other. Off-diagonal entries represent interspecific competition, while the diagonal entries represent self-regulation via intraspecific competition.

The second and third sub-matrices, *M_LT_* and *M_TL_*, describe the antagonistic relation between trees and lianas, that is the positive effect of the tree species on the lianas (providing physical support needed to reach the canopy (Burnham, 2002)) in combination with the negative impact of the lianas on the trees ((Estrada-Villegas et al., 2022; Estrada-Villegas and Schnitzer, 2018; García León et al., 2018; Meunier, 2026; Pérez-Salicrup, 2001; Schnitzer and Bongers, 2002; Schnitzer et al., 2005; Stevens, 1987; Stewart and Schnitzer, 2017)). Therefore, *M_LT_* contains only positive matrix elements, while *M_TL_*contains only negative matrix elements. In contrast to *M_TT_*, which is unstructured, *M_TL_* and *M_LT_* are assumed to have an internal architecture that reflects liana host preference, meaning that not all lianas can grow on all trees, as explained in subsection 2.2. The structure of *M_LT_* is mirrored in the structure of *M_TL_*, so that *M_LT__ij_* ≠ 0 if and only if *M_TL__ji_* ≠ 0.

The last sub-matrix, *M_LL_*, finally describes the interactions between liana species. We assume that lianas only interact with other lianas that share at least one host tree. The structure of *M_TL_* and *M_LT_*thus determines the position of non-zero elements in *M_LL_*. Whether or not these elements are then positive or negative depends on additional model assumptions. On the one hand, one can assume that lianas also compete for above-ground and below-ground resources, similar to the trees. However, the literature shows that lianas also benefit from the presence of other lianas, because they facilitate climbing on trees by climbing on one another (Ofosu-Bamfo et al., 2019; Pérez-Salicrup et al., 2001). Both positive and negative liana-liana interactions are thus conceivable. We therefore systematically vary the type of liana-liana interactions and distinguish between four cases: only positive matrix elements (translating into liana-liana facilitation, as shown in Fig 2), only negative matrix elements (translating into liana-liana competition, not shown) and two mixed cases (also not shown). In the first mixed case, each pair of lianas is assigned either positive or negative matrix elements (translating into pairwise facilitation or competition), whereas in the second mixed case matrix elements are completely independent from each other and can be either positive or negative (translating into liana-liana facilitation, competition or even antagonism).

In summary, our networks are thus conceptually similar to so-called multiplex networks, that is networks composed of multiple layers, each corresponding to a different interaction type (Boccaletti et al., 2014; Menichetti et al., 2014), which have received increasing attention in the context of ecology (see for example (Hervías-Parejo et al., 2020; Kéfi et al., 2016, 2017; Pilosof et al., 2017; Stella et al., 2018)). Assuming that the first layer would describe competitive interactions, the second layer antagonistic interactions and the third layer facilitative interactions, our community matrices could be translated into multiplex networks with either 2 or 3 layers, dependent on the type of liana-liana intraguild interaction.

### 2.2 Manipulating modularity in the antagonistic subnetwork

Modularity is a network property that measures the strength of a network’s division into subnetworks (also called modules; Alcalá-Corona et al. (2021); Dormann and Strauss (2014)). A highly modular network implies a pronounced difference between the connectance within modules and between modules, whereas no modularity implies the same connectance and thus no division into subnetworks (Alcalá-Corona et al., 2021; Fortunato, 2010; Newman, 2006). Assuming that all modules within the antagonistic sub-matrices are of the same size, we generate the antagonistic tree-liana interaction networks based on four parameters, namely the number of species *n_species_* = *T* = *L*, the connectance of the antagonistic part of the network *C*, the number of modules *n_m_*, and what we call the input modularity *m*. Given these parameters, we first determine the number of links inside and outside of predefined modules. The precise location of these links is then chosen at random.

The extreme case of maximum input modularity, *m* = 1, meaning that the antagonistic network can be divided into disjunct compartments, is relatively simple: all interactions are inside the predefined modules so that the number of links within modules equals *n_intra_*_,1_ = *C* · (*n_species_*)^2^ and the number of links outside of modules equals *n_inter_*_,1_ = 0 (see Fig. 2 a). Along similar lines, when modularity is absent, *m* = 0, meaning that all links are randomly distributed in the antagonistic sub-matrix, the number of links within modules equals 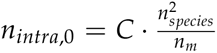 and the number of links outside of modules equals 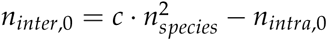.

Intermediate values of *m* translate into imperfect division into compartments (see Fig. 2 b). The number of interactions is linearly redistributed between the inter-and intra-module sets of possible interactions in proportion to the desired value of the input modularity *m*. The number of links within modules is then given by

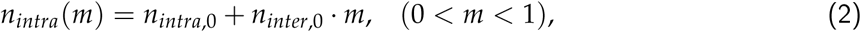

and the number of links outside of modules is given by

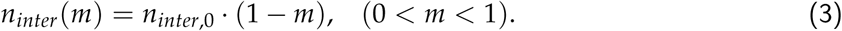

To justify this definition of modularity, we test whether varying *m* actually yields different levels of observed modularity. More precisely, we generate an ensemble of matrices with different values of *m* and measure the modularity of the resulting antagonistic subnetwork *M_TL_* via the Beckett algorithm, which is standard for computing the modularity of weighted bipartite networks in ecology (Beckett, 2016; Ofosu-Bamfo et al., 2022). We then look at the relation between the measured Beckett modularity, *Q*_Beckett_, and the input modularity, *m*, as shown in Fig. 3. We find that *Q*_Beckett_ increases along with *m* (Fig. 3), enabling us to generate networks with a measured modularity ranging between approximately 0.45 and 0.85.

**Figure 3:**
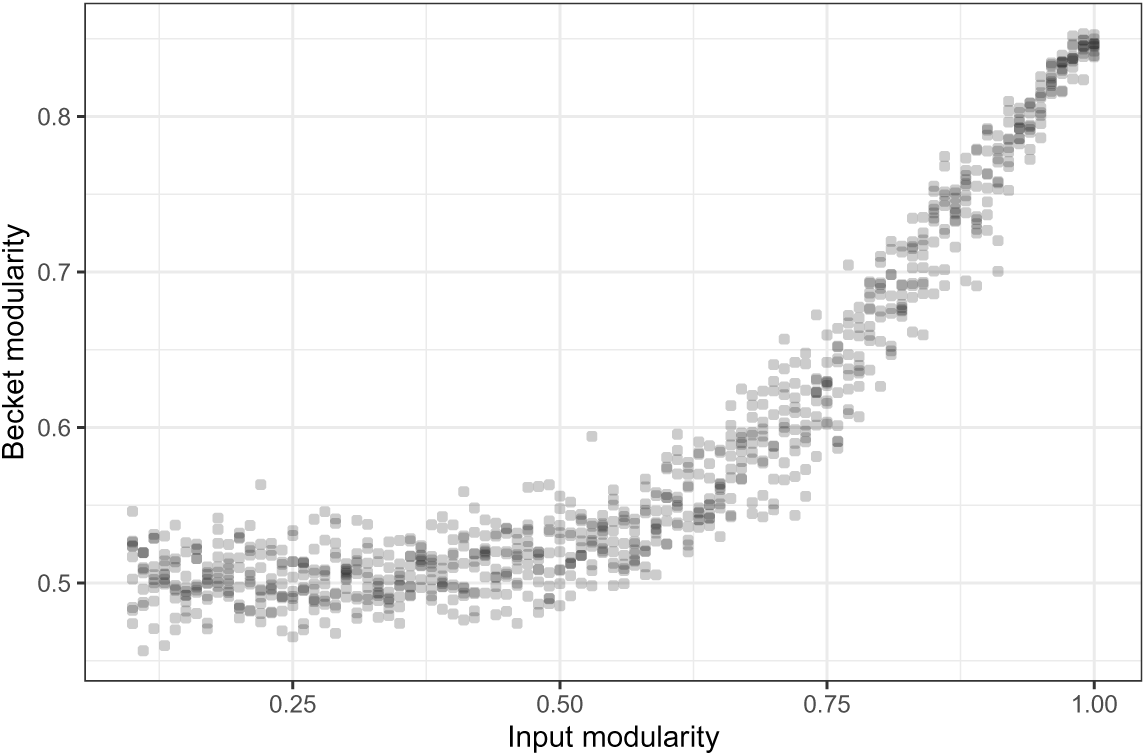
Relation between input modularity. *m* **and Beckett modularity**, using the LPAwbplus algorithm from the bipartite package, as introduced by Beckett (2016) and used in (Ofosu-Bamfo et al., 2022).

### 2.3 Assigning inter-and intraspecific interaction strengths

The structural properties that we vary in this study are defined by the locations of positive and negative interactions within the matrix, not by their absolute strengths. Still, to evaluate stability, we need to assign magnitudes to all non-zero matrix elements, representing the strength of those positive and negative interactions.

We set all diagonal matrix elements, which represent the strengths of self-regulation of each species to −1, following May (1972) and Allesina and Tang (2012). This assumes that our matrix elements are normalised interaction strengths, meaning that all off-diagonal elements represent the strength of interspecific interactions relative to the strength of intraspecific interactions in the matrix. The interaction strengths are thus dimensionless, which makes it possible to compare the stability of two matrices based on their eigenvalues (Neutel and Thorne, 2014; Thorne et al., 2021; Zelnik et al., 2024).

We furthermore choose the off-diagonal elements by drawing random numbers from a half-normal distribution. A half-normal distribution allows us to generate only positive or negative interactions depending on the interaction type and to easily transition to a normal distribution in the mixed-interaction case. We choose the same distribution, with a common scale parameter *σ* for all sub-matrices to make sure that the effect on stability is determined by modularity and interaction type, not by variation in interaction strengths.

Note that we focus on relative differences in stability and how these vary with modularity and intra-guild interaction type, not on the absolute values of *Re*(*λ_d_*). Negative values of *Re*(*λ_d_*) can be generated by increasing the strength of intra-relative to interspecific interactions, or in the case of normalised matrices, by decreasing the strength of interspecific interactions. Whether our community matrices are stable or not is thus an artefact of our parameter choice. As we have currently no empirical data available that would allow us to derive realistic estimates of interaction strengths, we arbitrarily chose a scale parameter of *σ* = 0.5 for our main analysis and performed additional robustness checks with varying *σ* values (see online Supplementary Material).

### 2.4 Feedback loop analysis

It has long been known that network stability is closely linked to the concept of feedback loops (Levins, 1974). A feedback loop is the effect of a species on itself via its impact on other species. For example, tree species A affects liana species B, which in turn affects tree species A, resulting in a closed chain of effects. In principle, feedback loops can be either positive (i.e., self-reinforcing), in which an initial perturbation is amplified, or negative (i.e., self-dampening), in which it is counteracted. Whether a feedback loop is positive or negative depends on the product of the interaction strengths involved. For example, the antagonistic feedback loop between tree species A and liana species B mentioned above is negative, because it involves one positive and one negative interaction. By contrast, a feedback loop between two tree species that compete with each other or two liana species that facilitate each other would be positive, resulting in self-reinforcing and hence destabilising feedback.

Whether a given system is stable generally depends on the complex interplay between all feedback loops within the system (Levins, 1974) but several studies that analysed feedback loops in ecological networks have shown that (in-)stability is mostly governed by relatively short feedback loops of length 2 and 3 (see (Neutel et al., 2002, 2007; Neutel and Thorne, 2014) for trophic networks and (Koch et al., 2026, 2023) for competitive networks). Therefore, we here also focus on loops of length 2 and 3. Note that in networks where structure is associated with patterns of strong and weak interaction strengths, it is important to compare the strength of feedback loops, which is determined by the product of all interaction strengths. In our case, however, as all interaction strengths are chosen independently from the same distribution, the hypothesised effect of modularity on stability must be driven via changes in the number of positive and negative loops, not their strength. We therefore simply count the number of positive and negative feedback loops of length 2 and 3, and classify these loops based on the species involved. For the latter, we distinguish between three different types of 2-link loops (liana-liana, tree-liana or tree-tree) and four different types of 3-link loops (liana-liana-liana, tree-liana-liana, tree-tree-liana and tree-tree-tree). We focus specifically on liana-liana, liana-liana-liana, tree-liana-liana and tree-tree-liana loops. We intentionally exclude tree-liana, tree-tree and tree-tree-tree loops from our analysis, since their number neither depends on the number of liana-liana interactions, nor on the degree distribution among the lianas, meaning that they are by design unaffected by varying the input modularity *m*.

### 2.5 Sensitivity analysis

If not stated otherwise, we used the standard parameter set given in Tab. 1. However, to ensure that our results do not depend on specific parameter choices or model assumptions, we additionally performed several robustness checks, as summarised in the online supplementary material. In particular, we tested whether the effect of modularity on system stability depends on the chosen system size, number of modules, connectance within the antagonistic sub-matrix or scale of the half-normal distribution from which we sample interspecific interaction strengths. Finally, we also tested whether model outcomes stay robust when using a gamma distribution instead of half-normal to assign interspecific interaction strengths.

## 3 Results

### 3.1 Modularity stabilises tree-liana networks

We analysed stability in an ensemble of community matrices with varying levels of modularity of the antagonistic tree-liana interactions and we repeated this analysis for facilitative, competitive as well as mixed liana intraguild interactions. We found that more modular antagonistic structures consistently yielded more stable communities (Fig. 4). We furthermore found that varying the type of liana intraguild interactions had a strong effect on stability in matrices with low levels of modularity, while for very high levels of modularity, all matrices showed roughly the same level of stability. Matrices with facilitative liana-liana interactions were most unstable when modularity was low and hence displayed a strong relationship between the level of modularity and stability. In contrast, matrices with competitive or mixed intraguild interactions were generally more stable and hence displayed a weaker stabilising effect of modularity. The weakest effect was observed in the mixed random case, including liana-liana facilitation, competition and antagonism. Our sensitivity analysis shows that these results are robust to parameter variation (smaller or larger networks, fewer or more modules, lower or higher connectance, less or more variation within interspecific interaction strengths) and also when using a gamma distribution instead of a half-normal distribution for generating interaction strengths, as summarised in the online supplementary material.

**Figure 4:**
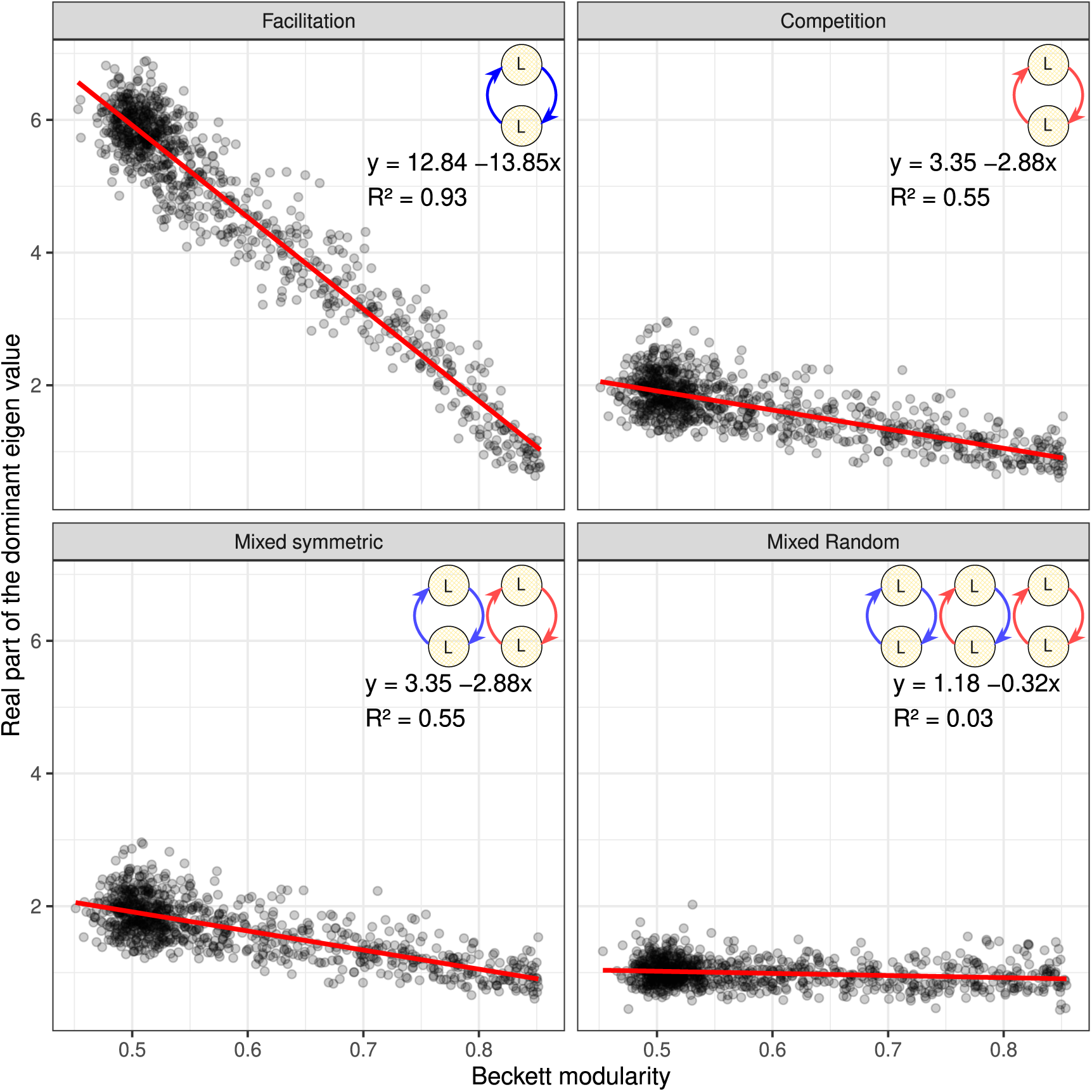
I**n**creasing **levels of modularity stabilise tree-liana networks**. Correlation between observed modularity and system stability for community matrices with liana-liana facilitation (top left), liana-liana competition (top right), mixed liana-liana interactions with positive and negative interactions coupled (bottom left) and mixed liana-liana interactions without structure (bottom right). We quantified system stability based on the real part of the dominant eigenvalue, with lower values indicating more stable systems.

### 3.2 Stabilising effect of modularity depends in intraguild interaction type

To better understand how variation in structure influences system stability, we analysed how increasing modularity influenced the number of positive and negative feedback loops in the system (Fig. 5). Higher levels of modularity reduce the connectance in the liana-liana sub-matrix, reflecting our assumptions that lianas only interact with other lianas sharing at least one host tree. Consequently, we found that a high level of modularity strongly reduced the number of liana-liana and liana-liana-liana feedback loops. The effect of modularity on the number of tree-liana-liana and tree-tree-liana loops was much weaker and the effect of modularity on the remaining loops (tree-tree, tree-liana and tree-tree-tree loops) was absent by design.

**Figure 5:**
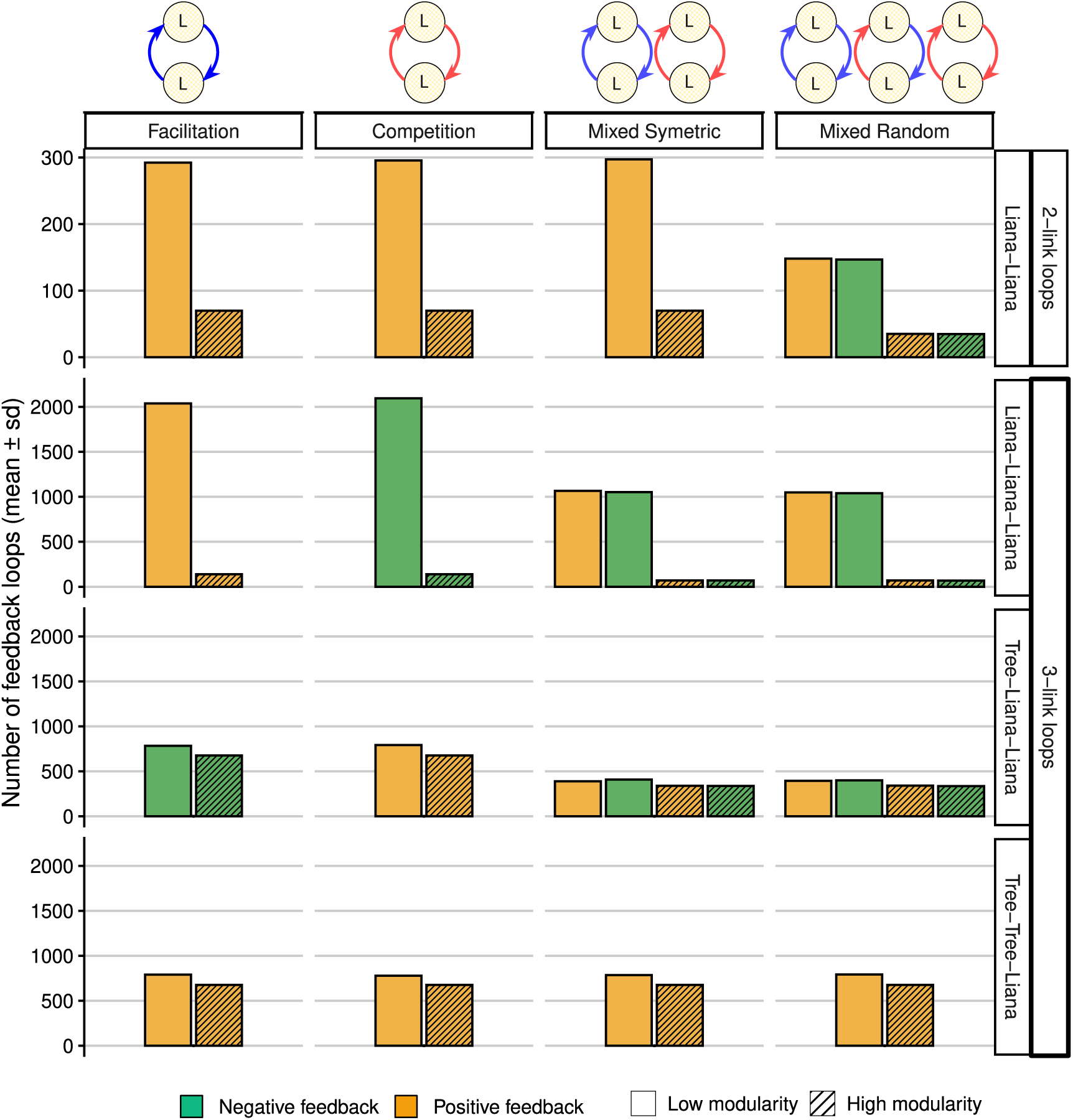
M**o**dularity **shifts the balance between positive and negative feedback loops.** Effect of modularity on the number of liana-liana, liana-liana-liana, tree-liana-liana and tree-tree-liana loops, assuming facilitative, competitive, mixed symmetric (facilitation and competition) or randomly mixed (faciliation, competition and antagonism) intraguild interactions. The number of tree-liana, tree-tree and tree-tree-tree loops is not shown because it is unaffected by varying the input modularity *m*.

Taken together, our feedback loop analysis suggests that the effect of modularity on stability was mostly driven by feedback loops formed by liana-liana interactions (Fig. 5). This explains why we found similar levels of stability in very modular matrices, regardless of intra-guild interaction types: For high modularity, feedback loops formed by liana-liana interactions are rarer and become negligible for overall system stability. In contrast, matrices with lower modularity contain many feedback loops formed by liana-liana interactions, explaining why the modularity-stability relationship depended strongly on the considered type of intraguild interaction (Fig 4).

In the case of liana-liana facilitation, both the 2-link liana-liana and the 3-link liana-liana-liana feedback loops are positive and hence create self-reinforcing feedback (Fig. 5). A reduction in positive 2-link loops then results in a stabilising effect of modularity, which is amplified by an additional reduction in positive 3-link loops, resulting in a relatively strong effect of modularity on stability. In contrast, in the case of liana-liana competition, we find that the 2-link liana-liana loops are positive, while the 3-link liana-liana-liana loops are negative. The stabilising effect of the reduction in positive 2-link loops is thus partly compensated by a reduction in negative 3-link feedback, resulting in a relatively weak effect of modularity on stability (Fig. 4).

When positive and negative interactions are mixed, either symmetrically or randomly, we find an equal number of positive and negative 3-link liana-liana-liana loops (Fig. 5). A reduction in the number of these loops hence does not translate directly into a stabilising or destabilising effect, so that the effect of modularity on stability is in these cases mostly driven by the reduction of 2-link loops. In the symmetric mixed case, we find that these 2-link loops are always positive, exactly as in the facilitation and the competition case, which explains why the effect of modularity on stability is weaker compared to the case of facilitation but stronger compared to the case of competition (Fig. 4). In contrast, in the random mixed case, we also find an equal number of positive and negative 2-link liana-liana loops (Fig. 5), which explains why the effect of modularity on stability is almost absent. The very small effect that we observed in Fig. 4 might be attributed to a small shift towards more negative feedback in the whole system: While the number of positive loops formed by liana-liana interactions decreased, negative feedback loops in other parts of the network remained unaffected (tree-liana, tree-tree-tree, not shown in the figure).

## 4 Discussion

In this study, we investigate the structure-stability relation of systems with multiple interaction types. More precisely, we consider systems that describe not only antagonistic interactions between two guilds (e.g., trees and lianas) but also competition, facilitation or even antagonism within each guild. We find that systems with a more modular arrangement of antagonistic interactions are generally more stable, a pattern that is independent of the specific parameter choice. We furthermore find that this stabilising effect of modularity strongly depends on the type of intraguild interaction under consideration. It is largest when intraguild interactions represent facilitation (e.g. lianas using already existing lianas to climb trees), intermediate when intraguild interactions represent competition (e.g. lianas competing for light accessibility and nutrients) or a mix of facilitation and competition and smallest when intraguild interactions are completely random. Finally, we find that all differences in effect sizes can be explained by a shift in the balance between self-reinforcing and self-damping feedback loops.

Our key result that modularity is stabilising is consistent with the results reported by Thébault and Fontaine (2010) and Stouffer and Bascompte (2011), indicating that this pattern can indeed be generalised from networks with a single interaction type to those with multiple interaction types. However, it is important to note that in our study, the stabilising effect of modularity is caused and modulated by feedback loops that emerge from intraguild interactions, which are neither considered in the bipartite networks studied by Thébault and Fontaine (2010) nor in the food webs studied by Stouffer and Bascompte (2011). This essential difference in model design suggests that the observed stabilising effect of modularity can arise from more than one mechanism. In our case, it is mostly caused by a reduction in destabilising 2-link and 3-link feedback loops, in particular when considering intraguild facilitation. The fact that the effect size of modularity varies for different intraguild interactions furthermore suggests that structure-stability relations in natural systems might change over time, for example when intraguild interactions are more facilitative in early successional stages of network development but more competitive in more mature communities (Malkinson et al., 2003; Woods et al., 2021), and also across space, for example due to gradients in density and/or environmental conditions (Holmgren et al., 1997; Zhang and Tielbörger, 2020). In the specific context of tree-liana systems, our results suggest that the effect of modularity on forest resilience is more pronounced at the highly disturbed forest edges, where positive co-occurrence of lianas on host trees was found, indicating liana-liana facilitation, whereas co-occurrence patterns were more random at interior, undisturbed habitats (Ofosu-Bamfo et al., 2022).

The insights gained from our study contribute to a growing body of literature investigating structure-stability relations in complex interaction networks containing different types of biotic interactions, as reviewed in Kéfi et al. (2017). Interestingly, many recent studies focus on tripartite systems with two different interaction types, such as plant-pollinator-herbivore or plant-herbivore-parasite systems (Domínguez-García and Kéfi, 2024; Emary and Malchow, 2022; García-Callejas et al., 2023; Sauve et al., 2014, 2016; Yan, 2022; Zhao et al., 2025). These systems essentially merge two bipartite webs by letting the species in one guild be the joining nodes between both networks. A common theme in these studies is then to show that the way these two bipartite networks are combined affects their overall stability. For example, Sauve et al. (2014) found that structures that promote stability in networks made of a single interaction type, such as nestedness and modularity, had weaker effects in networks merging mutualistic and antagonistic interactions. Several other network studies that also consider a mix of interaction types focus less on structural network properties and more on the question how shifting proportions of different link types affect system stability and species persistence (García-Callejas et al., 2018; Kondoh and Mougi, 2015; Lurgi et al., 2016; Mougi and Kondoh, 2012; Sellman et al., 2016; Suweis et al., 2014), leading to conflicting results. For example, García-Callejas et al. (2018) found that positive interactions are key for maintaining species persistence, whereas Sellman et al. (2016) reported a higher frequency of functional extinctions in networks with mixed interactions. We emphasise that our study is conceptually different from these two approaches, as we consider two guilds, similar to bipartite systems, but with additional intraguild interactions. To the best of our knowledge, the role of such intraguild interactions for structure-stability relations has received little attention (but see Yan (2022) for the effect of intraguild competition on the relation between nestedness and stability and García-Callejas et al. (2023) for the effect of interactions within and across guilds on coexistence patterns), which highlights the need to study how patterns in inter-and intraguild interactions jointly affect ecosystem stability.

Our study was inspired by Ghanaian tree-liana interaction networks (Addo-Fordjour and Afram, 2021; Addo-Fordjour et al., 2023; Ofosu-Bamfo et al., 2022) and might have important implications for their management and conservation. Within these ecosystems, both trees and lianas are key structural elements. Trees influence the forest structure by modulating light availability, nutrient cycling, and micro-climate regulation (Bongers et al., 2005). Lianas, on the other hand, contribute to the structure and function of tropical forests by providing fleshy fruits to frugivores (Estrada-Villegas and Schnitzer, 2018; Magrach et al., 2016), offering a dense micro-habitat for foraging and nesting, and serving as locomotion pathways for arboreal species such as the orangutan (Estrada-Villegas and Schnitzer, 2018; Magrach et al., 2016). Both trees and lianas are subject to harvesting, due to their economic and cultural value (Bongers et al., 2002; Cunningham et al., 2021), which raises important questions about how such interventions affect forest structure and resilience and which management strategies might be most suitable to achieve a balance between harvest yield on the one hand and ecosystem functioning on the other. In this context, our theoretical results suggest that management strategies that maintain modular network structures might be more sustainable than others, because they preserve a system’s ability to quickly return to equilibrium.

However, it is important to note that community matrices are very abstract and simplified representations of real communities and that further research is needed to better understand the real life consequences of network structure in tree-liana systems. Community matrices, as used in this study, allow us to obtain general insights into how modularity and intra-guild interaction type affected system stability and to test whether a community represented by a given matrix would return to steady state or not after a small perturbation, but they can not provide insights into actual trajectories of real world system after more realistic perturbations. To explore these dynamics in more detail, alternative approaches like individual-based models could be useful. Additionally, our results point towards the key role of liana intra-guild interactions for system stability, but these interactions have so far not been quantified in empirical studies. Furthermore, evidence for modular tree-liana network structures from locations outside Ghana is mixed (see for example Sfair et al. (2010)). Linking theoretical and empirical research will thus be necessary to fully understand the drivers of structural properties in observed tree-liana interaction networks.

The underlying assumptions in our model are very general and might apply not only to tree-liana networks, but to a broad range of other systems. This obviously includes systems with similar network structures, that is systems characterised by antagonistic interactions between two guilds in combination with various types of intraguild interactions, such as predator-prey systems with predator interference (Mougi, 2022) or host-parasite systems with competition or facilitation between parasites (Cusumano et al., 2016; Rodgers and Bolnick, 2024; Vaumourin et al., 2015). However, our results might also be of relevance for systems without inter-guild antagonism. In particular, we assume that the stabilising effects of nestedness within mutualistic communities, as found in single-interaction type networks (Okuyama and Holland, 2008; Rohr et al., 2014; Thébault and Fontaine, 2010), might be modulated, amplified or dampened when it appears within more complex networks with different types of intraguild interactions. In this context, our results highlight the need to develop a more holistic view on ecological networks, taking into account both inter-and intra-specific interactions, with the goal to develop a comprehensive understanding of structure-stability relations across ecosystems.

## Acknowledgments

We thank the whole department of Eco-Evolutionary Modelling and in particular Felix Jäger for insightful discussions and helpful feedback on the manuscript.

## Author contribution statement

RLD: Conceptualisation, Formal analysis, Investigation, Methodology, Software, Visualisation, Writing – original draft, Writing – review and editing

FK: Conceptualisation, Investigation, Methodology, Supervision, Writing – original draft, Writing – review and editing

BOB: Conceptualisation, Validation, Writing – review and editing

KTA: Conceptualisation, Funding acquisition, Investigation, Methodology, Project administration, Supervision, Writing – original draft, Writing – review and editing

