## Supplementary Material for "Stabilising effect of modularity in antagonistic networks depends on intraguild interactions"

### Supporting Information for "Stabilising effect of modularity in antagonistic networks depends on intraguild interactions"

To assess the robustness of our main results, we repeated the simulations under alternative parametrisations of network structure and interaction strengths. Specifically, we varied the number of species, the number of modules, the connectance within the antagonistic sub-matrix and the scale of the half-normal distribution from which we sample random interaction strengths. We also tested whether our results are robust when using a gamma distribution instead of a half-normal distribution. Across all tested scenarios, the qualitative relationship between modularity and network stability remained unchanged: higher modularity was generally associated with lower system instability, as summarised in Table S1. Additionally, also the overall pattern of feedback loop spectra was preserved (not shown here), with a consistent reduction in liana-liana and liana-liana-liana feedbacks across simulations. However, the strength of the relationship between modularity and stability varied across parameter settings, as explained in more detail below, indicating that some network configurations were more sensitive to changes in modularity than others.

#### S1 Number of species

Increasing species richness generally increased the explanatory power of modularity, whereas low species richness yielded substantially weaker fits. This reduction in predictive power at low species richness may reflect the smaller number of possible interactions and hence the greater sensitivity of small systems to stochastic variation.

#### S2 Number of modules

Changing the number of modules had only a limited effect on the estimated slopes and coefficients of determination. This can be explained by the fact that the number of modules detected by the Beckett algorithm differs from the input number of modules, since it uses a clustering algorithm. Thus, the variation in detected modules is less drastic than that in input modules.

#### S3 Connectance of antagonistic sub-matrix

Connectance had a stronger effect on the fit in the competition scenario, where high connectance was associated with a marked reduction in the coefficient of determination. This result likely reflects the narrower range of modularity values observed under this parametrisation, which limits the variation available to detect a relationship between modularity and stability. For the other interaction types, the effect of connectance on model fit was weaker.

#### S4 Scale of the half-normal distribution

For the half-normal distribution, varying the scale parameter had only a minor effect on the coefficient of determination, although larger variance produced generally more unstable matrices (see section 2.3 in the main text) and thus higher intercept values and steeper slopes.

#### S5 Gamma distribution

There is little literature on the statistical distribution of interaction strengths, as measuring them in the field is very challenging (Wootton and Emmerson, 2005; Novella-Fernandez et al., 2019). However, we know that ecological networks typically contain many weak and only a few strong links, resulting in link distributions that are highly skewed (Berlow, 1999; O'Gorman et al., 2010; Jacquet et al., 2016; Wootton and Emmerson, 2005; Paine, 1992). In fact, this pattern has been found in systems as diverse as bryozoans (Koch et al., 2026), algae (Paine, 1992), and plant-pollinator networks (Jordano, 1987). The half-normal distribution that is typically used in theoretical studies (including ours) does not reflect this high level of skewness. We therefore tested whether our results remain robust when using a gamma distribution. We found that the results were overall very similar compared to results obtained with the half-normal distribution.

Table S1: List of the robustness checks. All parameters of the simulations are set to the standard values as in Table 1 of the main manuscript, except for the parameter indicated in the left column. The slope and the intercept of the regression indicate the magnitude of the effect of modularity on stability. The value of $R^2$ (coefficient of determination) indicates the proportion of variation in stability that is explained by variation in modularity.

| | | Slope | Intercept | $R^2$ | Lowest Beckett modularity | Highest Beckett modularity |
|---|---|---|---|---|---|---|
| Reference scenario (see Fig 4 in the main manuscript) | Competition | -0.87 | 1.61 | 0.24 | 0.45 | 0.85 |
| | Facilitation | -14.08 | 12.96 | 0.94 | | |
| | Mixed symmetric | -2.86 | 3.36 | 0.55 | | |
| | Mixed random | -0.43 | 1.23 | 0.06 | | |
| High number of species Number species = 40 | Competition | -0.97 | 1.81 | 0.3 | 0.45 | 0.85 |
| | Facilitation | -16.15 | 15.26 | 0.95 | | |
| | Mixed symmetric | -3.65 | 4.66 | 0.74 | | |
| | Mixed random | -0.46 | 1.42 | 0.07 | | |
| Low number of species Number species = 15 | Competition | -0.44 | 0.57 | 0.01 | 0.55 | 0.85 |
| | Facilitation | -0.97 | 1.02 | 0.05 | | |
| | Mixed symmetric | -0.43 | 0.6 | 0.01 | | |
| | Mixed random | -0.22 | 0.37 | 0 | | |
| High number of modules Number modules = 8 | Competition | -0.85 | 1.58 | 0.12 | 0.45 | 0.75 |
| | Facilitation | -12.56 | 12.07 | 0.81 | | |
| | Mixed symmetric | -2.99 | 3.43 | 0.34 | | |
| | Mixed random | -0.27 | 1.14 | 0.01 | | |
| Low number of modules Number modules = 3 | Competition | -1.83 | 2.08 | 0.2 | 0.45 | 0.7 |
| | Facilitation | -18.08 | 14.91 | 0.78 | | |
| | Mixed symmetric | -4.03 | 3.95 | 0.28 | | |
| | Mixed random | -0.67 | 1.36 | 0.03 | | |
| High connectance Connectance = 0.18 | Competition | -0.48 | 1.4 | 0.01 | 0.45 | 0.625 |
| | Facilitation | -12.49 | 12.09 | 0.46 | | |
| | Mixed symmetric | -3.23 | 3.57 | 0.1 | | |
| | Mixed random | -0.85 | 1.45 | 0.02 | | |
| Low connectance Connectance = 0.08 | Competition | -0.75 | 1.52 | 0.2 | 0.45 | 0.85 |
| | Facilitation | -13.33 | 12.45 | 0.93 | | |
| | Mixed symmetric | -2.91 | 3.4 | 0.57 | | |
| | Mixed random | -0.43 | 1.23 | 0 | | |
| Half-normal high scale $\sigma = 2$ | Competition | -3.4 | 9.33 | 0.22 | 0.45 | 0.85 |
| | Facilitation | -54.95 | 53.89 | 0.93 | | |
| | Mixed symmetric | -11.57 | 16.47 | 0.56 | | |
| | Mixed random | -1.64 | 7.91 | 0.05 | | |
| Half-normal low scale $\sigma = 0.1$ | Competition | -0.18 | -0.48 | 0.24 | 0.45 | 0.85 |
| | Facilitation | -2.76 | 1.76 | 0.93 | | |
| | Mixed symmetric | -0.6 | 0.11 | 0.57 | | |
| | Mixed random | -0.09 | -0.55 | 0.06 | | |
| Gamma same mean and variance as standard $\alpha = 1.75$, $\theta = 0.228$ | Competition | -0.8 | 1.56 | 0.2 | 0.45 | 0.85 |
| | Facilitation | -13.57 | 12.6 | 0.93 | | |
| | Mixed symmetric | -2.79 | 3.32 | 0.55 | | |
| | Mixed random | -0.33 | 1.17 | 0.04 | | |
